# Lost in translation: egg transcriptome reveals molecular signature to predict developmental success and novel maternal-effect genes

**DOI:** 10.1101/286815

**Authors:** Caroline T. Cheung, Thaovi Nguyen, Aurélie Le Cam, Amélie Patinote, Laurent Journot, Christelle Reynes, Julien Bobe

## Abstract

**Background:** Good quality or developmentally competent eggs result in high survival of progeny. Previous research has shed light on factors that determine egg quality, however, large gaps remain. Initial development of the embryo relies on maternally-inherited molecules, such as transcripts, deposited in the egg, thus, they would likely reflect egg quality. We performed transcriptome analysis on zebrafish fertilized eggs of different quality from unrelated, wildtype couples to obtain a global portrait of the egg transcriptome to determine its association with developmental competence and to identify new candidate maternal-effect genes.

**Results:** Fifteen of the most differentially expressed genes (DEGs) were validated by quantitative real-time PCR. Gene ontology analysis showed that enriched terms included ribosomes and translation. In addition, statistical modeling using partial least squares regression and genetics algorithm also demonstrated that gene signatures from the transcriptomic data can be used to predict reproductive success. Among the validated DEGs, *otulina* and *slc29a1a* were found to be increased in good quality eggs and to be predominantly localized in the ovaries. CRISPR/Cas9 knockout mutants of each gene revealed remarkable subfertility whereby the majority of their embryos were unfertilizable. The Wnt pathway appeared to be dysregulated in the *otulina* knockout-derived eggs.

**Conclusions:** Our novel findings suggested that even in varying quality of eggs due to heterogeneous causes from unrelated wildtype couples, gene signatures exist in the egg transcriptome, which can be used to predict developmental competence. Further, transcriptomic profiling revealed two new potential maternal-effect genes that have essential roles in vertebrate reproduction.

## Background

Good quality or developmentally competent fish eggs are defined as those that are successfully fertilized and develop normally as viable, non-malformed embryos that hatch[1]. However, the detailed mechanisms that are involved in egg quality and developmental competence are still poorly understood, and at present, no predictive markers of egg quality exist. Maternal-effect genes are those that produce factors that are involved in the earliest stages of embryonic development, including fertilization, parental genome union, and cell division. Since initial development of the embryo relies on these maternally-inherited molecules including coding and non-coding mRNAs and proteins that are deposited into the developing oocyte, thus, they would likely reflect egg quality[2,3]. Among these, the maternally-provided transcriptome of the egg is critical in kick-starting early embryogenesis because transcription from the zygotic genome does not start until the mid-blastula transition (MBT) which occurs approximately 3-4 hours post-fertilization (hpf) in zebrafish[4,5].

Previous research using both traditional mutational assays as well as more recent transcriptomic analyses have revealed several maternal factors that can influence egg quality. The nucleoplasmin 2 (*npm2a* and *npm2b*) genes were recently found to be crucial for egg developmental competence; suppression of *npm2b* resulted in embryonic arrest before zygotic genome activation (ZGA) in mouse and zebrafish, and *npm2a* deficiency in zebrafish led to a complete lack of embryonic development[6]. Further, post-ovulatory ageing induced egg quality defects are associated with low mRNA levels of *igf1* (insulin growth factor 1) and beta-tubulin, as well as a small but significant overabundance of keratins 8 and 18, cathepsin Z, and *pgs2* (prostaglandin synthase 2)[7,8]. In addition, controlled induction of ovulation by hormonal or photoperiod manipulation negatively impacts egg quality in rainbow trout, and the abundance of several genes including *apoC1* (apolipoprotein C1), *mr-1* (major histocompatibility class 1 related protein), *ntan1* (N-terminal asparagine amidase 1), *myo1b* (myosin 1b), *pyc* (pyruvate carboxylase), as well as *phb2* (prohibitin 2) was found to be significantly different between naturally and artificially induced eggs[9]. Other studies have suggested that genes involved in immune regulation have an impact on egg competence whereby variable abundance of transcripts in the interferon pathway and *mhc* (major histocompatibility) class genes was demonstrated in eggs of different quality[10,11]. However, despite these results, knowledge on the factors that contribute to the quality of fish eggs remains patchy. Thus, in this study, we carried out a large-scale analysis to compare the transcriptome of eggs of different quality and performed statistical modeling of differentially expressed genes (DEGs) with survival in order to determine if there are common factors that impact egg quality in unrelated wildtype (WT) females that can then serve as markers and/or predictors of developmental competence. We further conducted functional analyses on two candidate genes that were increased in bad quality eggs using the CRISPR/cas9 knockout system and reveal for the first time the essential roles of two new potential maternal-effect genes, *otulina* (OTU deubiquitinase with linear linkage specificity a) and *slc29a1a* (solute carrier family 29, member 1a). Our findings provide evidence that even in different quality eggs from unrelated, wildtype couples bred under standard conditions, gene signatures exist in the egg transcriptome, which can be used to predict developmental competence, and that two new potential maternal-effect genes have essential roles in vertebrate reproduction.

## Results

### Transcriptomic differences between good and bad eggs in all samples

Among the 136 clutches of fertilized egg we collected, we selected 16 clutches each of good and bad quality eggs defined as those with >93% and <38% survival at 48 hpf, respectively, for microarray analysis using a customized chip containing 61,657 annotated sequences of the zebrafish transcriptome. We excluded the sequences of which 80% of the samples did not have any expression from further analyses, and we identified 31,261 annotated sequences that were expressed in the majority of the samples. Using the GeneSpring software with an FDR <0.05 as an exclusion criteria, 66 DEGs that were statistically significant between good and bad quality eggs were revealed. We observed in the heat map showing supervised clustering (Fig. 1a) of the 66 DEGs that a majority of them were upregulated (60 genes, red signal) with only a few genes (6 genes) that were decreased (blue signal) in bad quality eggs as compared to good quality eggs. Additional file 1 lists the 66 DEGs including their associated information. Of these 66 genes, 8 were annotated in Ensembl with a unique identifier, but were not found to be associated with any known gene or protein.

**Fig. 1 a:**
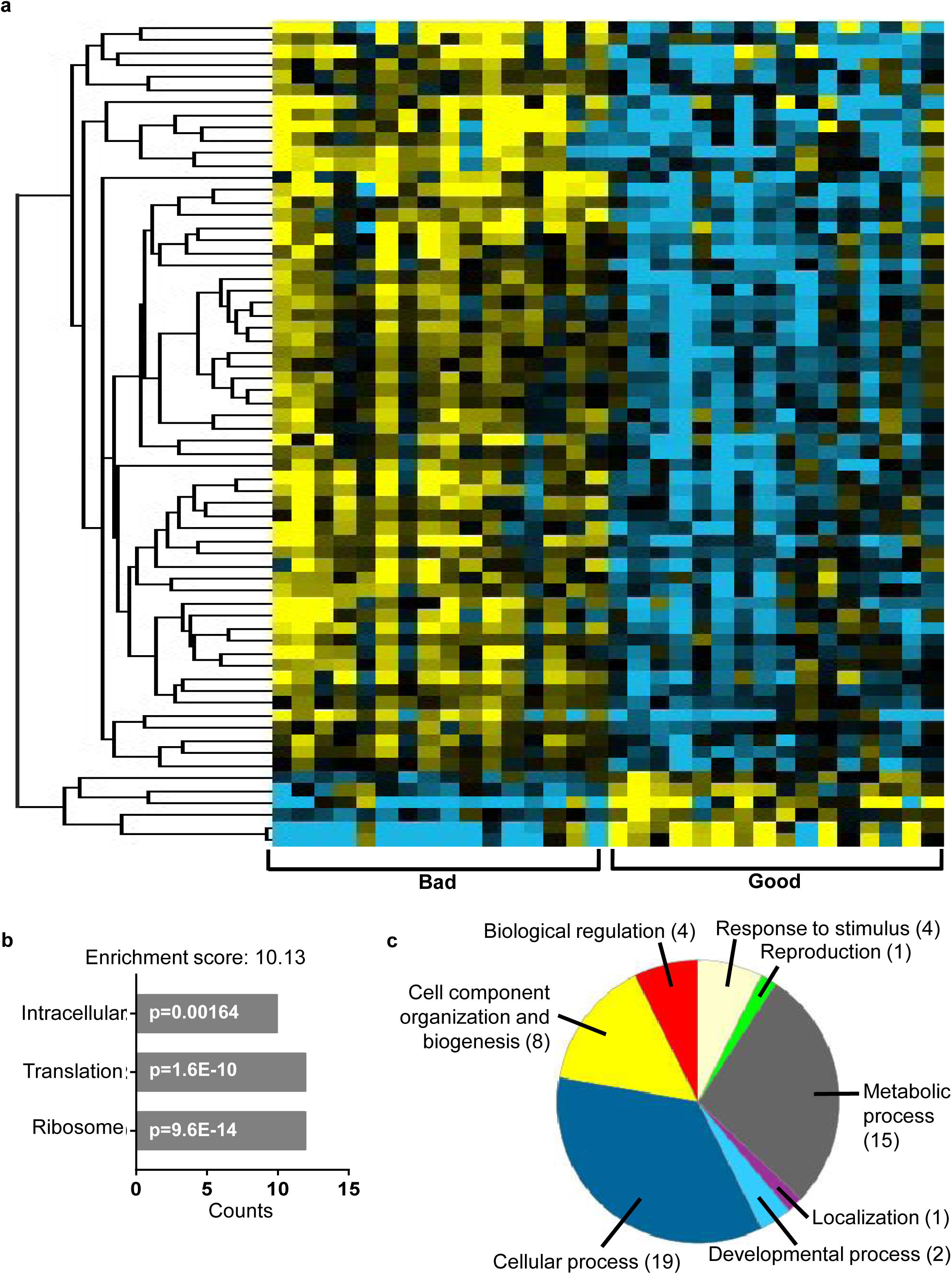
Heat map showing supervised clustering of the 66 differentially expressed genes (DEGs) between good and bad quality eggs from 32 clutches of fertilized zebrafish eggs. Yellow signal denotes upregulation, blue signal denotes downregulation, and black defines no change in expression. b: Gene ontology analysis using the DAVID online program of the 55 DEGs with known information. The enriched term and Benjamini value are shown. c: Functional classification of 54 DEGs with known function using the PANTHER online program. The biological processes associated with the DEGs and Benjamini value are shown. A Benjamini value of less than 0.05 was considered as statistically significant.

### Overrepresentation analyses of gene ontological terms of the DEGs

We submitted the 66 DEGs to functional annotation analyses by gene ontology using two different online programs, DAVID[12] and PANTHER[13], with the entire zebrafish transcriptome as background (Fig. 1b-1c). Among the 66 DEGs submitted, functional terms associated with 55 and 54 annotated genes were identified and therefore classified by the DAVID and PANTHER programs, respectively. In the former analysis, one cluster of terms (enrichment score: 10.13) were enriched from our gene list included intracellular (p=0.00164), translation (p=1.6E-10), and ribosomes (p=9.6E-14) (Fig. 1b). In the latter analysis, the DEGs were classified according to biological processes, which included cellular processes, metabolic processes, cell component and biogenesis, biological regulation, response to stimulus, developmental processes, localization, and reproduction (Fig. 1c). In fact, upon inspection of the differentially expressed genes shown in Additional file 1, the ones that underwent the most drastic changes in expression (ribosome production factor 2 homolog (*S. cerevisiae*) [*rpf2*], ribosomal protein S27 (isoform 2) [*rps27*], and U1 spliceosomal RNA [*U1*] with fold changes of 7.81, 1.90, and -2.33/-2.35, respectively) are associated with translation//ribosomes.

### Quantitative real-time polymerase chain reaction (qPCR) validation of the DEGs

In order to confirm the results obtained by microarray analysis, another independent method to detect gene expression changes was performed. qPCR was conducted using the same 32 samples that were submitted to microarray analysis and the primers used are listed in Additional file 2. Eight genes that underwent the most drastic changes in microarray analysis were subjected to qPCR, and their biological function as well as the p-value and fold change in the microarray analysis are shown in Additional file 3. qPCR confirmed that the expression of *rpf2* (1.87±0.33 vs. 0.48±0.20, p=0.01), *spon1b* [spondin 1b] (1.61±0.34 vs. 0.49±0.09, p=0.0003), *tspan7b* [tetraspanin 7b] (1.00±0.11 vs. 0.50±0.08, p=0.001), *rps27* (2.82±0.18 vs. 1.66±0.13, p<0.0001), *stra13* [stimulated by retinoic acid 13 homolog/centromere protein X] (1.20±0.09 vs. 0.87±0.12, p=0.03), and *rtn4ip* [reticulon 4 interacting protein 1] (1.02±0.07 vs. 0.84±0.04, p=0.03) were upregulated in bad quality eggs as compared to good quality eggs, while that of *U1* (21.08±5.81 vs. 4.38±1.28, p=0.009) and *slc29a1a* (1.04±0.05 vs. 1.26±0.06, p=0.008) were increased in bad relative to good quality eggs (Fig. 2a-h). Interestingly, despite the statistical significance in the differential regulation of *U1* (Fig. 2g), the expression of this gene was regulated in the opposite direction by qPCR as compared to by microarray analysis. In fact, we found that *U1* expression was decreased on average by 2.3-fold in bad quality eggs relative to good quality eggs as assessed by microarray, but qPCR results showed that it was increased by approximately 5-folds in bad as compared to good quality eggs. Regardless of this difference, we found by both microarray and qPCR that the transcript level of all eight genes were differentially regulated.

**Fig. 2:**
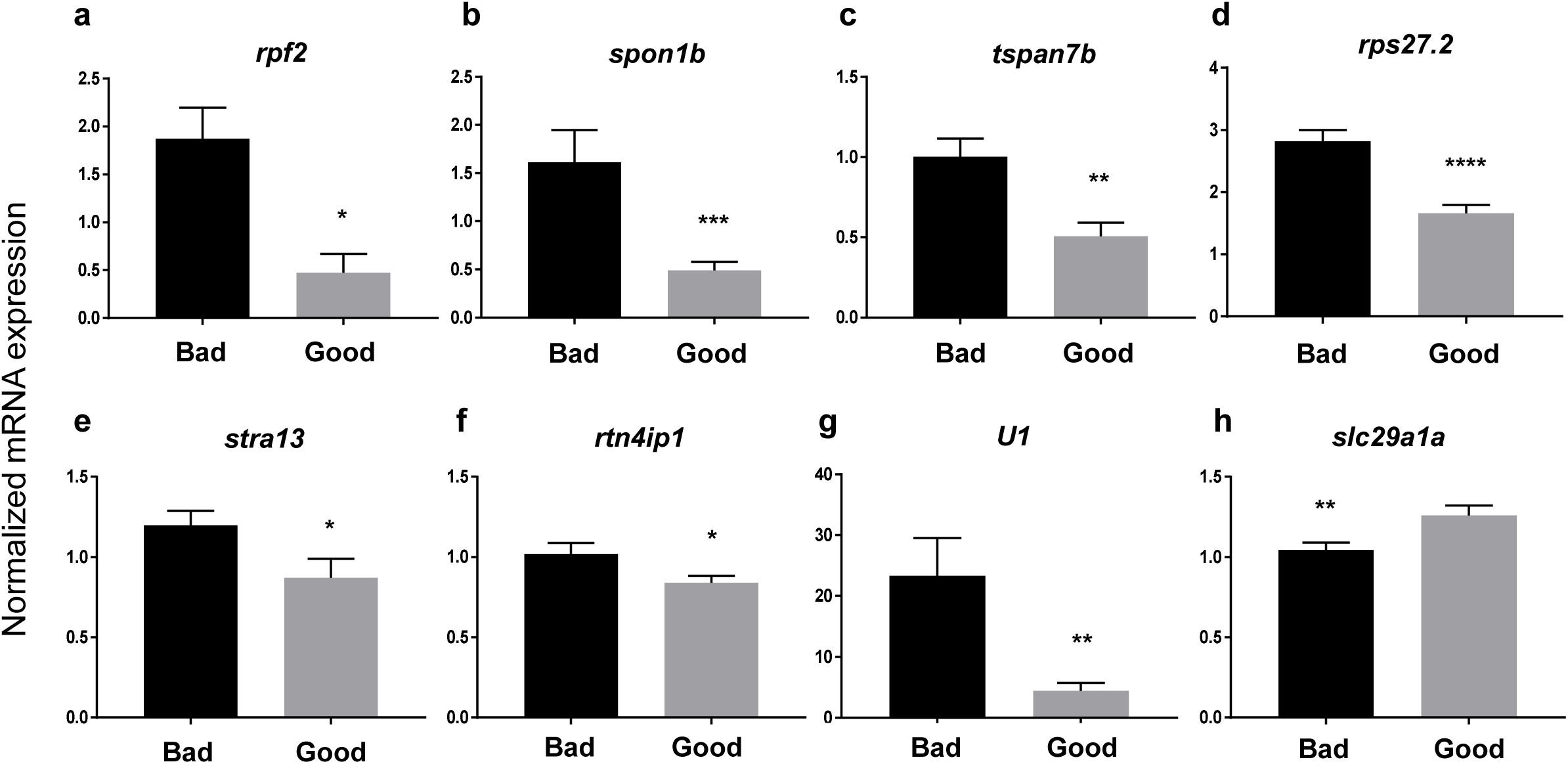
Validation of the microarray data by performance of quantitative real-time PCR (qPCR). Eight genes, including (a) *rpf2*, (b) *spon1b*, (c) *tspan7b*, (d) *rps27.2*, (e) *stra13*, (f) *rtn4ip1*, (g) *U1*, and (h) *slc29a1a* were subjected to qPCR using the primers listed in Additional file 2, whereby LSM couples member 14B (*lsm14b*), prefoldin subunit 2 (*pfdn2*), and ring finger protein 8 (*rnf8*) as well as 18S rRNA, *beta-actin* (*bact*), and *elongation factor 1 alpha* (*EF1α*) were used as internal controls. * p-value ≤0.05, ** p-value ≤0.01, *** p-value ≤0.001, **** p-value≪0.001.

### Couples analysis

Within the two groups of fertilized eggs, we observed a large variability in the expression of the genes using both detection methods. Upon further inspection, we found that certain couples (#5, 10, 33; Additional file 4) consistently produced bad quality eggs (≤50% survival at 48 hpf). Therefore, fertilized eggs harvested at two different periods (1-3 months apart) from these 3 couples (6 samples in total) along with samples from 6 random couples that consistently produced good quality eggs were submitted for re-analyses by microarray and qPCR.

### Transcriptomic differences by microarray in different couples

Microarray analysis using the GeneSpring program of the six samples that came from the three couples that consistently produced bad quality eggs revealed 1385 DEGs, and there appeared less variability in transcript levels within each sample group, as shown in the heat map by supervised clustering (Fig. 3a). The complete list of annotated genes is shown in Additional file 5. Similar to the microarray results of all 32 samples, we observed that there were far more upregulated genes (1240) in the bad quality eggs relative to good quality eggs (145 genes). Of the 1385 DEGs, 233 could be identified with an Ensembl annotation, but were not found to be associated with any characterized gene or protein. We also observed that the differences in alterations in the transcript levels were much greater in the couples analysis than those detected when all samples were included. For example, there was a 53-fold increase in *tk2* [thymidine kinase 2], 25-fold upregulation in *drd3* [dopamine receptor D3], and a 60-fold decrease in the expression of *prkcq* [protein kinase C, theta] (Additional file 6), while the most differentially regulated genes (*rpf2, spond1b, tspan7b*, and *U1*) were altered by only 2-7-folds in the former analysis (Additional file 3).

**Fig. 3 a:**
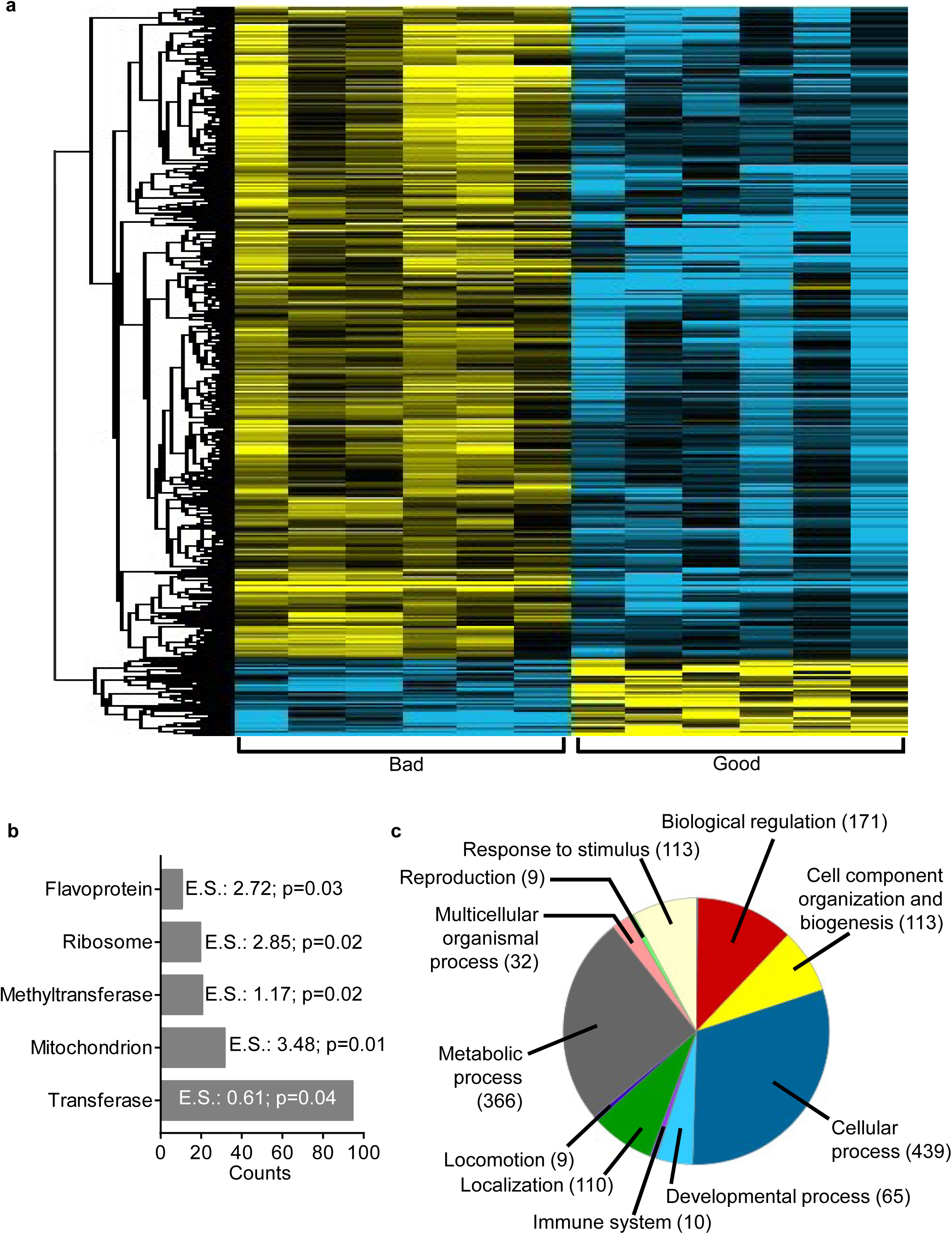
Heat map showing supervised clustering of the 1385 differentially expressed genes (DEGs) between good and bad quality eggs from 16 clutches of fertilized zebrafish eggs in the couples analysis. Red signal denotes upregulation, blue signal denotes downregulation, and black defines no change in expression. b: Gene ontology analysis of the 1151 DEGs with known information using the DAVID online program. The enriched term and Benjamini value are shown. c: Functional classification of the 1135 DEGs with known function using the PANTHER online program. The biological processes associated with the DEGs and Benjamini value are shown. A Benjamini value of less than 0.05 was considered as statistically significant.

### Overrepresentation analyses of gene ontological terms of the DEGs in the couples analysis

In order to get an idea of the functional properties of the 1385 DEGs found by the couples analysis, we submitted this list of genes for gene ontology as before. Of the 1385 DEGs, we were able to classify and identify functional properties of 1151 and 1135 of the genes by the DAVID and PANTHER programs, respectively. The DAVID analysis revealed 5 clusters with enriched terms including ribosome/translation (enrichment score: 2.85; p=0.02) as well as mitochondria (enrichment score: 3.48; p=0.01), flavoprotein (enrichment score: 2.72; p=0.03), methyltransferase (enrichment score: 1.17; p=0.02), and transferase (enrichment score: 0.61; p=0.04) (Fig. 3b). The PANTHER analysis revealed that the DEGs were involved in many very similar biological processes as the previous analysis such as cellular processes, metabolic processes, biological regulation, cell component and biogenesis, response to stimulus, localization, developmental processes, multicellular organismal process, immune system, locomotion, and reproduction. Ribosome/translation appeared to be the term that was greatly enriched in all the analyses, suggesting that genes that function in this process are especially important in determining egg quality.

### qPCR validation of the DEGs in the couples analysis

qPCR was performed to validate the dysregulation of the DEGs that were modified the most in the couples analysis, as listed in Additional file 6 along with their known biological process as well as the p-value and fold change as assessed by microarray. *rpf2* and *tspan7b* were also found to be drastically dysregulated in the couples analysis, thus, they were resubmitted for qPCR using just the 12 samples. We confirmed by qPCR that the transcript levels of *tk2* (0.97±0.41 vs. 0.0002±0.0001, p=0.004), *drd3* (3.04±0.32 vs. 0.25±0.09, p=0.0022), *rpf2* (2.82±0.28 vs. 0.58±0.39, p=0.009), *cldn23* [claudin 23] (4.43±0.42 vs. 0.82±0.58, p=0.004), *tspan7b* (1.11±0.11 vs. 0.46±0.15, p=0.03), and *stra13* (1.47±0.17 vs. 0.61±0.08, p=0.0022) were increased (Fig. 4a-f), while *pomt1* [protein-O-mannosyltransferase 1] (0.37±0.04 vs. 1.37±0.20, p=0.0022), *prkcq* (0.003±0.001 vs. 6.02±1.67, p=0.0022), *nudt13* [nucleoside diphosphate linked moiety X-type 13] (0.02±0.01 vs. 1.82±0.60, p=0.0022), *itih2* [inter-alpha-tryspin inhibitor heavy chain 2] (0.05±0.02 vs. 1.42±0.31, p=0.0022), *flvcr1* [feline leukemia subgroup C cellular receptor family, member 2a] (0.05±0.01 vs. 3.07±1.02, p=0.0022), *otulina* (0.10±0.02 vs. 0.53±0.07, p=0.0001), and *slc29a1a* (1.04±0.04 vs. 1.36±0.13, p=0.05) were downregulated in bad relative to good quality eggs in these 12 samples (Fig. 4g-m).

**Fig. 4:**
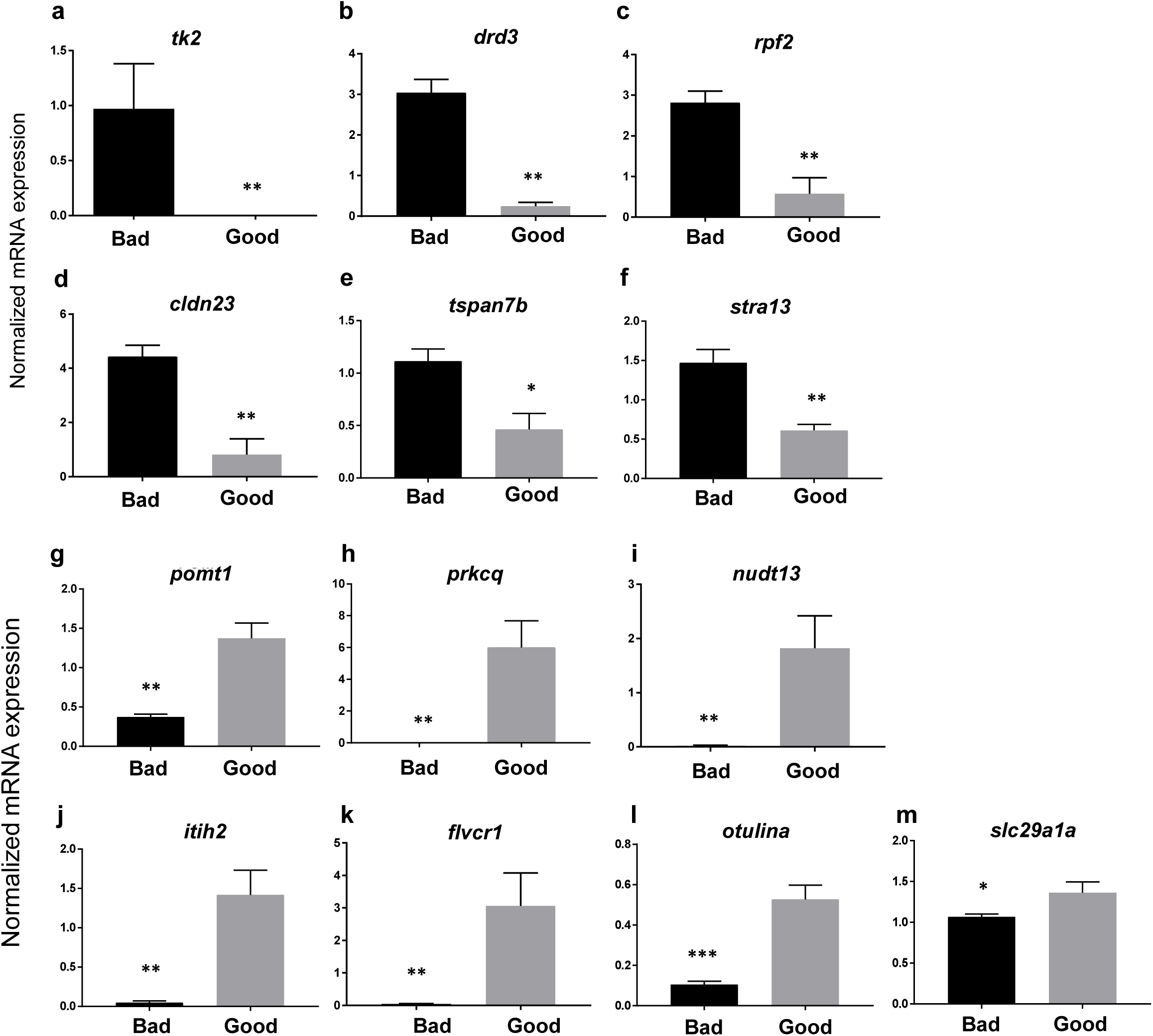
Validation of the microarray data by performance of quantitative real-time PCR (qPCR). The six genes upregulated, (a) *tk2*, (b) *drd3*, (c) *rpf2*, (d) *cldn23*, (e) *tspan7b*, (f) *stra13*, and seven genes downregulated, (g) *pomt1*, (h) *prkcq*, (i) *nudt13*, (j) *itih2*, (k) *flvcr1*, (l) *otulina*,(m) *slc29a1a*, in bad quality eggs were subjected to qPCR using the primers listed in Additional file 2. LSM couples member 14B (*lsm14b*), prefoldin subunit 2 (*pfdn2*), and ring finger protein 8 (*rnfB*) as well as 185 rRNA, beta-act n (*bact*), and *elongation factor 1 alpha* (*EF1α*) were used as internal controls. * p-value ≤0.05, ** p-value ≤0.01, *** p-value ≤0.001

### Functional analysis of otulina and slc29a1a in zebrafish

In order to validate the *in vivo* significance of some of the DEGs, we performed functional analysis by genetic knockout using the CRISPR/cas9 system on *otulina* and *slc29a1a*. RNA-seq data stored in the PhyloFish[14] online database (Additional file 7) as well as qPCR analysis for *otulina* (Fig. 5a) and *slc29a1a (*Fig. 5b) in different tissues revealed that both of these genes were expressed predominantly in the ovary, which suggest that they play a role in oogenesis and/or reproduction and are thus good candidates for knockdown. One-cell staged embryos were injected with the CRISPR/cas9 guides that targeted either *otulina* or *slc29a1a* and allowed to grow to adulthood. Mosaic founder mutant females (F0) were identified by fin clip genotyping and subsequently mated with wild-type (WT) or Dr_*vasa:*eGFP C3 (hereafter called *vasa*:eGFP) males, and embryonic development of the F1 fertilized eggs was recorded. Since the mutagenesis efficiency of the CRISPR/cas9 system was very high, as previously described[15,16], the *otulina* and *slc29a1a* genes were sufficiently knocked-out even in the transgenic mosaic F0 females. This was evidenced by the substantially lower transcript levels of *otulina* and *slc29a1a* in the F1 embryos as compared to those from control WT pairings (Fig. 5c). Thus, the phenotypes of *otulina* (n=4) and *slc29a1a* (n=10) mutants could be observed even in the F0 generation. Since none of the mutated genes were transmissible to future generations neither through the male nor the female (ie. only WT F1 progeny survived until adulthood), therefore, all of our observations were obtained from the F0 generation.

**Fig. 5:**
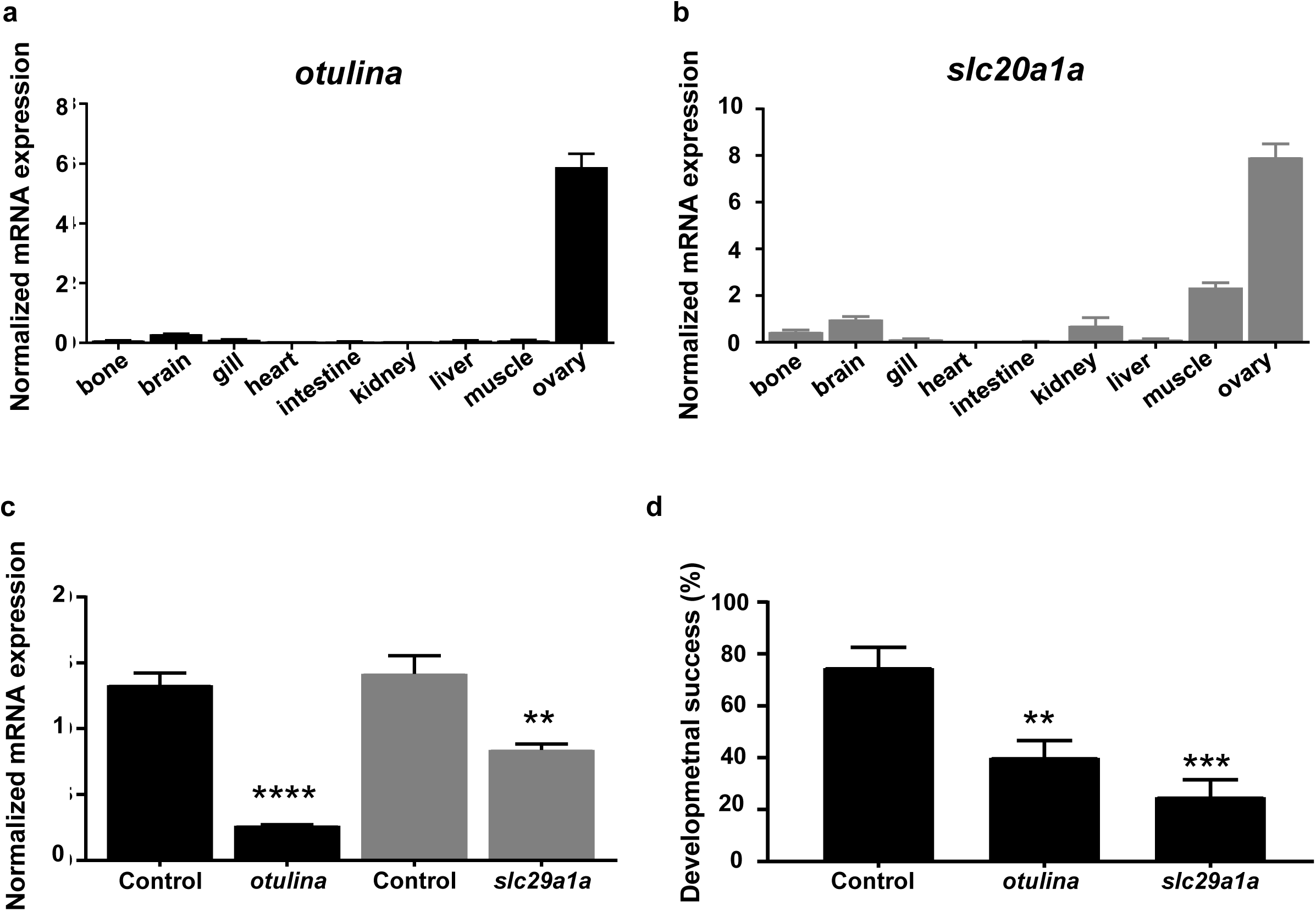
Tissue localization of *otulina* (a) and *slc29a1a* (b) based on qPCR assays. c: Expression level of *otulina* and *slc29a1a* in spawned eggs from mutant females mated with WT males as assessed by qPCR. 18S rRNA, *beta-actin* (*bact*), and *elongation factor 1 alpha* (*EF1α*) were used as internal controls, and experiments performed in triplicate. d: Developmental success in terms of survival rate of embryos at 24 hours post-fertilization (hpf) from *otulina*- and *slc29a1a*-deficient mutant females mated with WT males. N=4 for *otulina* and N=10 for *slc29a1a*, at least three spawns from each mutant.

We observed that both *otulina* and *slc29a1a* mutant-derived eggs had a very low developmental success, defined as the proportion of surviving embryos at 24 hpf to the total number of spawned eggs (40.0±6.7% and 24.8±6.8%, respectively, vs 74.61±7.9% in controls) (Fig. 5d). One spawn of fertilized eggs from the cross between each individual mutant female and a *vasa:eGFP* male was counted based on its developmental phenotype, described as non-cellularized (lack of cell division), partially cellularized (abnormal cell division), and normal development, as shown in Table 1. As compared to the control embryos that developed normally from 2-24 hpf (Fig. 6a-d), most of the spawned eggs from the mutant females were non-cellularized such that they did not undergo any cell division at all throughout the same time period, and they eventually all died by 24 hpf (Fig. 6e-l). However, two of the *slc29a1a* mutants displayed some heterogeneity in their offspring; while a proportion of the spawn did not develop and did not undergo cell division as observed previously, a number of cells underwent abnormal development characterized by asymmetrical cell division and the appearance of a cell mound on top of an enlarged cytoplasm, which occurred until approximately 4-5 hpf (Fig. 6m and 6n), after which they began to develop normally albeit slightly slower than their control counterparts (Fig. 6o-6q). To determine if the non-cellularized eggs were unfertilized or were arrested in development immediately after fertilization, we performed PCR genotyping for the *gfp* gene, which would only come from the *vasa:eGFP* male and not from the mutant mother that does not harbour any *gfp gene*. We found that the non-cellularized eggs from both *otulina* and *slc29a1a* mutants did not have the *gfp* gene indicating that they were not fertilized (Fig.6R). These novel findings showed for the first time that *otulina* and *slc29a1a* are essential for the developmental competence of eggs, and are therefore crucial maternal-effect genes.

**Table 1.** (please refer to pg. 10): Penetrance of *fam105ba* and *slc29a1a* mutant phenotypes.

|  | Total<br>number<br>of<br>embryos | Embryos with defects |  | Normal<br>embryos |
| --- | --- | --- | --- | --- |
|  |  | Non-cellularized | Partially<br>cellularized |  |
| *fam105ba-1* | 210 | 163 |  | 47 |
| *fam105ba-2* | 213 | 172 |  | 41 |
| *fam105ba-3* | 92 | 53 |  | 39 |
| *fam105ba-4* | 116 | 110 |  | 6 |
| *slc29a1a-1* | 104 | 104 |  | 0 |
| *slc29a1a-2* | 100 | 88 |  | 12 |
| *slc29a1a-3* | 660 | 450 |  | 210 |
| *slc29a1a-4* | 451 | 439 |  | 12 |
| *slc29a1a-5* | 245 | 150 |  | 95 |
| *slc29a1a-6* | 152 | 138 |  | 14 |
| *slc29a1a-7* | 361 | 252 |  | 110 |
| *slc29a1a-8* | 80 | 71 |  | 9 |
| *slc29a1a-9* | 85 | 9 | 48 | 28 |
| *slc29a1a-10* | 370 | 153 | 24 | 193 |

**Fig. 6:**
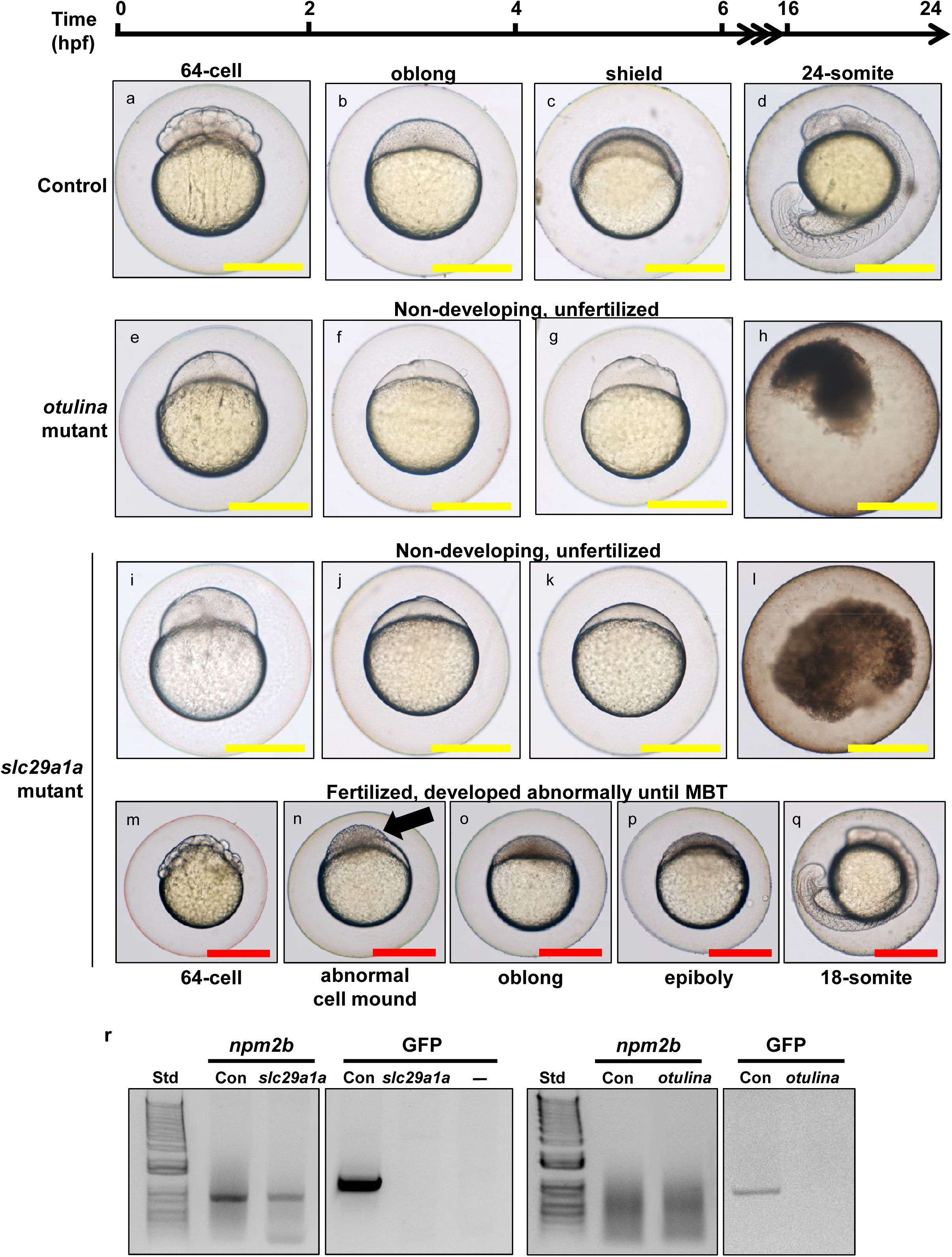
Representative images showing the development between 0-24 hours post-fertilization (hpf) of F1 embryos from wildtype control (a-d), *otulina*-deficient (e-h), and *slc29a1a*-deficient (i-q) females. In the control eggs, the embryos were at 64-cell (a), oblong (b), shield (c), and 24-somite (d) stages according to Kimmel et al[64]. Eggs from *otulina* and *slc29a1a* mutant females were non-developing and did not under any cell division (e-l). Some eggs from two *slc29a1a* mutant females were developing abnormally (m-q). (a, e, i, m) = images taken at 2 hpf; (b, f, j, n) = images taken at 4 hpf; (c, g, k, o) = images taken at 6 hpf; (p) = image taken at 8 hpf; (d, h, l, q) = images taken at 24 hpf. The arrow demonstrates a partially cellularized blastodisc that was sitting atop an enlarged syncytium. Scale bars denote 500 μm. r: PCR genotyping for *nucleoplasmin 2b* (*npm2b*) and *vasa*:*eGFP* in spawned eggs from WT, *otulina-*, and *slc29a1a*-mutant females crossed with *vasa:eGFP* males to detect fertilization of the eggs. Std = 1 kb ladder; Con = WT female crossed with *vasa:eGFP* male.

### The Wnt pathway is dysregulated following otulina deficiency

In a bid to elucidate a possible mechanism that may govern the function of *otulina*, we assessed the spawned eggs from *otulina*-mutant females crossed with WT males for the transcript levels of components of the *wnt* (*wnt3a, tcf3, tcf7, ef1*, and *dvl2*) and *tnf*/*nf*-*κb* (*nf-kb2, rel, rela, ikkaa, ikkab*, and *tnfa*) pathways. Previous studies have shown that *otulina* plays a role in these pathways in mammalian models[17–19]. Our findings showed that *wnt3a, tcf7, lef1*, and *dvl2*, but not *tcf3*, were significantly decreased in the *otulina* mutant-derived eggs (Fig. 7a-d), while none of the transcripts belonging to the *tnf*/*nf*-*κb* pathways were changed (Additional file 8). Thus, these results highly suggest that *otulina* plays a role in the *wnt* pathway in early development, and *otulina* deficiency leads to loss of developmental success due to dysregulation of *wnt* signaling.

**Fig. 7:**
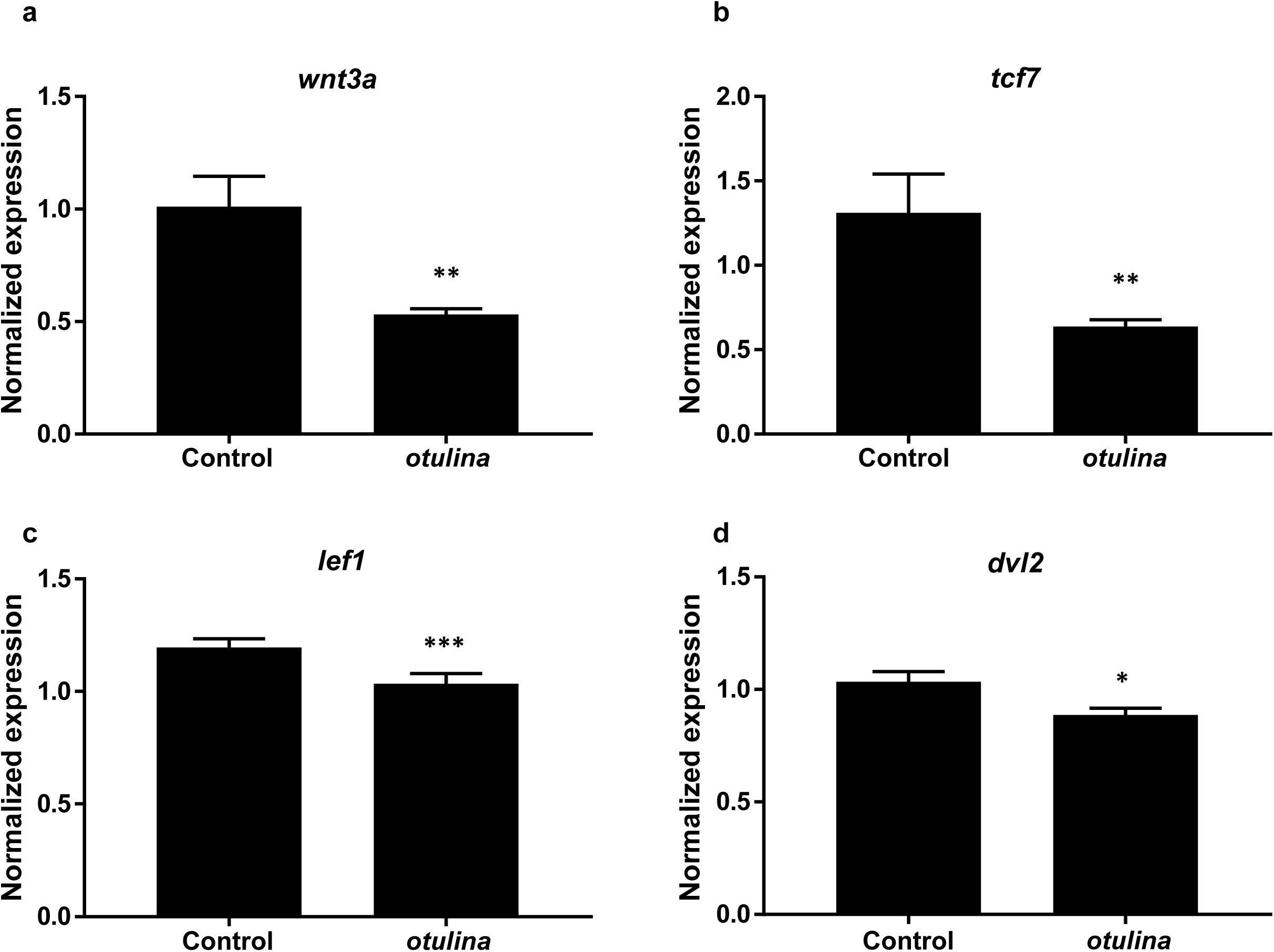
Evaluation of the expression levels of *wnt3a* (a), *tcf7* (b), *lef1* (c), and *dvl2* (d) in spawned eggs from *otulina*-deficient mutant females mated with WT males as assessed by qPCR. 18S rRNA, *beta-actin* (*bact*), and *elongation factor 1 alpha* (*EF1α*) were used as internal controls, and experiments performed in triplicate. N=4, at least three spawns from each mutant.

### Identification of gene signatures to predict developmental competence by statistical modeling using Partial Least Square (PLS) regression and genetic algorithm

We demonstrated in the previous figures that some of the differentially regulated genes were correlated with survival. However, these findings were based on univariate analysis of individual genes. Thus, we decided to use PLS regression to model the link between transcriptomic data and survival rate. Then, in order to select the best subsets of genes (of both small size and high survival prediction ability), a genetic algorithm was used as it allows the efficient exploration of sets of solutions (here subsets of genes) that are too abundant to be exhaustively explored.

In order to make the selection by genetic algorithm easier and quicker, a preliminary filtering of genes was performed: a correlation test performed on all 31,317 genes eliminated those that did not show any correlation at all to survival, and the redundant genes with highly correlated expressions were grouped together, which left us with 5410 genes for further analysis. Genetic algorithm coupled with PLS was applied to the 5410 remaining genes to search for subsets of genes potentially associated with survival. Two outputs can be expected from this analysis: first by globally exploring the solutions of several runs, it is possible to provide a list of genes which can be considered as potentially related to survival and, second, a more specific study of the solutions of several runs will allow the retrieval of a few candidate signatures of small subsets of complementary genes that we can use for diagnostic purposes.

We ran the genetic algorithm coupled with PLS 70 times with populations of 500 individuals. We thus obtained 35,000 final solutions. In order to set a frequency threshold to decide which genes can be selected, pseudodata were used: the genetic algorithm was applied using identical conditions as previously mentioned to datasets with randomly permuted survival rates. The 35,000 solutions were evaluated by 10 runs of 2-fold cross validation (2-FCV), and the average R^2^ values obtained from the 10 runs were retained as quality criteria for each solution. Hence, to ensure the proposed solutions were relevant, the average 2-FCV R^2^ values obtained on the actual dataset were compared to the ones obtained on the pseudo-datasets with permuted survival rates. Fig. 8a shows that the 2-FCV R^2^ obtained for the final solutions for the actual survival rates were significantly higher than that for the randomized ones (p-value <2.10E-16; Mann-Whitney U-test), thus, the prediction models are relevant. Further, the more often a gene appeared in those final solutions, the more likely it was associated with survival. Hence, if a variable was selected more often in the results obtained from the actual survival rates than from the pseudo survival rates, it may be considered to be potentially related to the phenomenon studied. We therefore compared how often each variable was selected in populations from the actual data and from the randomized data. The results are presented in Fig. 8b. The upper panel displays the selection frequencies of each gene in the final populations of the genetic algorithm applied on the real dataset while the lower panel displays those for the randomized values. These data clearly show that there were no significant peaks in the randomized data as compared to the actual dataset, which suggest that they do not carry biological meaning and can be used as a minimum frequency threshold for gene selection to be considered as biologically relevant. The 95th and 99th percentiles of the distribution of frequencies in the randomized data were used to obtain sets of genes that were the most often selected; 156 genes were obtained with a threshold of 5% and 29 genes with a threshold of 1%, the latter of which is listed in Table 2. Among these 29 genes, 10 of them were DEGs found by our microarray analyses (boldfaced, italicized).

**Table 2.** (please refer to pg. 13): List of 29 genes that are associated with survival.

| ENSEMBL gene annotation | Gene name |
| --- | --- |
| ENSDARG00000090871 | Si:dkey-210j14.4 |
| ENSDARG00000076419 | Si:dkeyp-117b11.2 |
| ENSDARG00000079255 | Zgc:174935 |
| ***ENSDARG00000031366*** | ***Reticulon 4 interacting protein 1*** |
| ENSDARG00000006982 | muscle segment homeobox D |
| ENSDARG00000071553 | Zgc:171500 |
| ENSDARG00000070898 / ENSDARG00000092291 | Si:ch211-262h13.3 / Si:ch211-281g2.3 |
| ENSDARG00000075318 | Solute carrier family 16 (monocarboxylic acid transporters), member 6a |
| ***ENSDARG00000063295*** | ***Myosin, heavy polypeptide 9a, non-muscle*** |
| ***ENSDARG00000082140 / ENSDARG00000082017*** | ***U1 spliceosomal RNA*** |
| ENSDARG00000089078 | Collagen, type XXIII, alpha 1 |
| ***ENSDARG00000017820*** | ***Polymerase (RNA) III (DNA directed) polypeptide D*** |
| ENSDARG00000024687 | Polymerase (RNA) III (DNA directed) polypeptide G |
| ***ENSDARG00000089422*** | ***CABZ01087562.1*** |
| ENSDARG00000056563 | Peroxisome proliferative activated receptor, gamma, coactivator 1, beta |
| ***ENSDARG00000076498*** | ***Golgi integral membrane protein 4a*** |
| ***ENSDARG00000069425*** | ***Heat shock factor binding protein 1a*** |
| ENSDARG00000090804 | G protein-coupled receptor 155a |
| ENSDARG00000075434 | RNA 2,,3,-cyclic phosphate and 5,-OH ligase |
| ENSDARG00000096436 | Si:dkey-118j18.4 |
| ENSDARG00000089677 | CABZ01117603.1 |
| ***ENSDARG00000095796*** | ***Si:dkey-87o1.2*** |
| ENSDARG00000004898 | Zona pellucida glycoprotein 2, like 2 |
| ENSDARG00000020149 | Acyl-Coenzyme A oxidase-like |
| ENSDARG00000027738 | Si:ch211-13c6.2 |
| ENSDARG00000080245 | 5S ribosomal RNA |
| ENSDARG00000058445 | Protein disulfide isomerase-like, testis expressed |
| ***ENSDARG00000078785*** | ***Transmembrane protein 258*** |
| ***ENSDARG00000093926 / ENSDARG00000095522*** | ***Si:dkey-71b5.2 / Si:dkey-71b5.3*** |

The 29 genes that were selected from the exhaustive analysis by Partial Least Square (PLS) regression and genetic algorithm. The 10 genes that were differentially regulated in our microarray dataset are boldfaced and italicized.

**Fig. 8 a:**
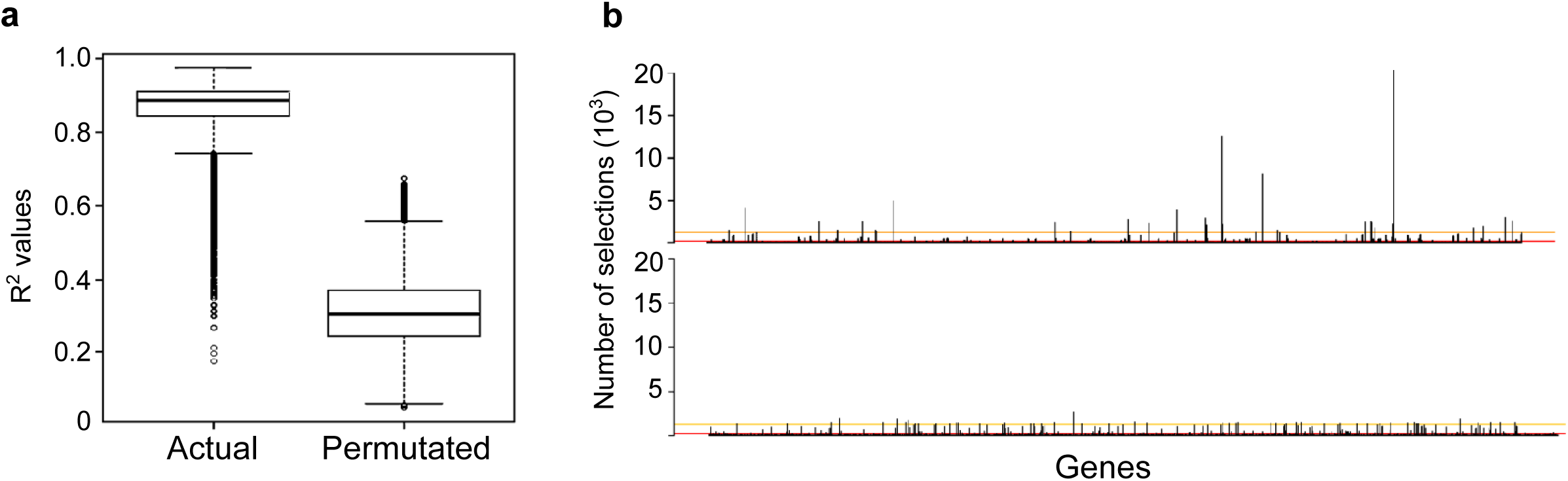
The average 2-fold cross validation R^2^ values obtained from the actual dataset were compared to the ones obtained from the pseudo-datasets with permuted survival rates. b: The frequency that each variable was selected in populations from the actual data and from the randomized data. The 95th and 99th percentiles of the distribution of frequencies in the randomized data were used to obtain sets of genes that were the most often selected.

Subsequently, the same results can be used to identify a gene signature that only retains small subsets of genes that are the most relevant for diagnostic purposes. Towards this goal, all 35000 final solutions were evaluated by 10 runs of 2-fold cross validation (2-FCV), and the average R^2^ values obtained from the 10 runs were retained as quality criteria for each solution. However, genetic algorithms are powerful methods that may be misleading, thus, the two best solutions were selected as good compromises between quality of the prediction and parsimony of the model (Table 3). The average 2-FCV R^2^ of solutions 1 and 2 were 0.9771 and 0.9678, respectively using 7 and 8 genes respectively (with 5 common genes between them, italicized).

**Table 3.** (please refer to pg. 13): Two solutions from the parsimonic prediction model.

| ENSEMBL gene ref | Gene name |
| --- | --- |
| **Solution 1** |  |
| *ENSDARG00000079255* | *Zgc:174935* |
| *ENSDARG00000089677* | *CABZ01117603.1* |
| *ENSDARG00000090871* | *Si:dkey-210j14.4* |
| *ENSDARG00000076419* | *Si:dkeyp-117b11.2* |
| *ENSDARG00000017820* | *polymerase (RNA) III (DNA directed) polypeptide D* |
| ENSDARG00000086485 | novel protein coding gene |
| ENSDARG00000020054 | aldehyde oxidase 1 |
| **Solution 2** |  |
| *ENSDARG00000079255* | *Zgc:174935* |
| *ENSDARG00000089677* | *CABZ01117603.1* |
| *ENSDARG00000090871* | *Si:dkey-210j14.4* |
| *ENSDARG00000076419* | *Si:dkeyp-117b11.2* |
| *ENSDARG00000017820* | *polymerase (RNA) III (DNA directed) polypeptide D* |
| ENSDARG00000016855 | Splicing factor 3b, subunit 5 |
| ENSDARG00000088305 | CABZ01072929.1 |
| ENSDARG00000087431 | Zgc:173962 |

Two solutions containing 7 and 8 genes that were selected from the parsimonic model by Partial Least Square (PLS) regression and genetic algorithm. The 5 common genes between the two solutions are italicized.

Thus, these findings demonstrated the presence of strong gene signatures, which were statistically robust both in terms of reproducibility and validation by pseudo-data, to link gene expression to the survival rate of eggs in our transcriptomic data.

## Discussion

In this study, we analyzed the maternally-provided transcriptome in fertilized zebrafish eggs at the one-cell stage in order to determine expressional differences between bad and good quality eggs that could impact egg quality in unrelated wildtype females. Despite the haphazard causes of decreased egg competence, we still found by microarray and validated by an independent method of transcript quantitation, qPCR, a large number of DEGs that were represented predominantly by those that function in ribosome and translation processes. Thus, there appears to be substantial differences in the maternally-inherited transcriptome between good and bad quality eggs that may have an impact on egg competence and subsequent embryonic development, and may serve as markers or predictors of egg quality.

When all the samples were taken into consideration, only 66 DEGs were found between good and bad quality eggs, which is a relatively low number as compared to other studies. In fact, it must be reiterated that all of the couples that were mated and produced clutches were wildtype, without any particular treatment, and mostly unrelated. Thus, there may have been multiple natural causes behind the decline in quality of the eggs from the different mothers, such as nutrition, density, age of parents, delay from last spawn, and genetics just to name a few[1,20,21]. Despite the heterogeneous potential causes of varying egg quality, 66 DEGs were identified of which 7 genes were verified independently by qPCR. These genes all play different cellular roles: *rpf2*[22] is a ribosome assembly protein that recruit 5S rRNA and ribosomal proteins into nascent large ribosome subunits; *rps27.2*[23] is a structural component of the 40S small ribosome subunit; *spond1b*[24] encodes a protein secreted by floor plate cells during embryogenesis that localizes to the central spinal canal and has neuroregulatory functions; *rtn4ip*[25] is a mitochondrial protein present in neurons and astrocytes; *tspan7b*[26] is a cell surface receptor signaling molecule that functions in embryonic development; *stra13*[27] has roles in DNA repair and kinetochore assembly; and *slc29a1a*[28] is transmembrane glycoprotein that mediates the cellular uptake of nucleosides. Their distinct roles in the cell highlight the fact that embryonic survival is based on many different cellular processes and suggest that they may serve as candidate markers of egg quality among unrelated wildtype females in larger populations.

A recent paper demonstrated that the transcriptome in unfertilized zebrafish eggs from different mothers can be quite variable while it was more uniform within clutches, which suggests that mother-related differences in the transcriptome may potentially be associated with egg quality and subsequent development of the embryo[29]. To this end, we performed a couples analysis in which we compared clutches of eggs from 3 couples that consistently produced bad quality eggs to those that frequently gave good quality eggs in order to eliminate some of the environmental effects on maternal health and egg competence. In this narrowed-down analysis using only 12 clutches of eggs (6 each of good and bad quality), we revealed more than 1300 DEGs with less variability between samples than in the first analysis (Figure 3). We also found that the 4 DEGs that were common in the two analyses, *rpf2, slc29a1a, tspan7b*, and *stra13* were more homogeneously expressed as there was less variability between samples. These results suggest that despite the heterogeneous causes, many common mechanisms contribute to egg quality among unrelated couples that consistently produce bad quality eggs, and these can serve as good markers for egg competence and hence embryonic survival.

With regard to the functional characteristics of the DEGs in the study using all samples, overrepresentation analyses of the GO terms by both DAVID and PANTHER programs found that genes that function in ribosome and translation were enriched, which are consistent with previous findings that showed that translation-related transcripts were also differentially expressed in sea bass eggs of different quality[30]. Interestingly, the findings in this study correlate with our previous proteomic study which also demonstrated significant dysregulation of proteins with functions in protein synthesis in zebrafish eggs of varying quality[31]. In the proteomic study, peptides that function in protein synthesis were upregulated in both good and bad quality eggs, which suggest a general dysregulation of the system. In this study, *rpf2* and *rps27.2* were found by microarray and confirmed by qPCR to be increased in bad quality eggs (Fig.2A, 4C, and 2D, respectively). Both of these genes encode proteins that function in ribosomes; rpf2 is an assembly factor and rps27.2 is a structural component of the 40S small ribosome subunit as mentioned above. A similar finding was demonstrated in a previous study that investigated the transcriptome of eggs after natural and controlled ovulation in rainbow trout (*Oncorhynchus mykiss*); it was revealed that *rpl24*, which is a structural component of the large ribosome subunit, was more abundant in the latter which had higher mortality[9]. Thus, it appears that in eggs of bad quality, there is an apparent increase in ribosome biogenesis. This seems somewhat contradictory due to the fact that most bad quality eggs eventually die and an accumulation in ribosomes is usually a sign of increased cell proliferation and growth[32]. It is well-known that ribosome biogenesis deficiency often leads to an impairment in cell growth and to cell death while elevated ribosomal function results in increased cell cycle progression and proliferation[33]. Thus, it is possible that the increased ribosomal content and presumably elevated translation that were found in bad quality eggs may reflect dysregulated cell cycle progression and cell growth, which in fact may lead to premature cell death as shown by pathological models in which loss of cell cycle and cell death regulators lead to disease[34,35]. On the other hand, it is also a possibility that ribosome biogenesis dysregulation may be an effect and not cause of loss of embryonic survival; an unknown mechanism that operates during oogenesis may cause the accumulation of ribosomal content in bad quality eggs and its abundance reflects a dysfunction of the mechanism. Whichever the case, dysregulation of the protein synthesis process appears to be at both the transcript and protein levels.

In general, the expression data in the couples analysis were less heterogeneous with less spread within each group, and the differences in gene expression between the two groups were larger than those observed in the first analysis (Fig. 4a-m and Additional file 6 vs. Fig. 2a-h and Additional file 3, respectively). Literature search showed that most of these genes have known functions: *tk2*[36] is a deoxyribonucleoside kinase that is required for mitochondrial DNA synthesis; *drd3*[37] is dopamine receptor which is associated with cognitive, emotional, and endocrine functions; *cldn23*[38] is an integral membrane protein that maintains cell polarity and signal transductions; *nudt13*[39] is hydrolase and its function is also largely unknown; *itih2*[40] is a serine protease inhibitor that carries hyaluronan in plasma; *flvcr1*[41] is a heme transporter that has a critical role in erythropoiesis by protecting developing erythroid cells from heme toxicity; *pomt1*[42] is an O-mannosyltransferase that is a component of many polysaccharides and glycoproteins; *prkcq*[43] is a serine- and threonine-specific protein kinase that is important for T-cell activation; and *otulina*[44] is a predicted deubiquitinase also called *fam105ba*. qPCR confirmed the differential expression of these genes in bad quality eggs relative to good quality eggs (Fig. 4a-m). From these results, it appears that there is a general dysfunction of multiple cellular processes when only couples that consistently produce bad quality eggs are taken into account, as opposed to a general population of random couples. Thus, the mechanisms that are involved in egg quality may be different depending on the couple; infrequent production of bad quality eggs in a random, unrelated population may involve dysfunction of ribosome/translation processes while there may be a general dysregulation of multiple cellular processes in frequent producers of bad quality eggs. Our results also suggest that within a random population of unrelated females, some common elements that impact egg competence exist, thus, a portfolio of gene profiles can be established for use as markers of egg quality which would be extremely useful in identification of reproductively successful females.

In order to determine if panels or sets of genes that together may be used to predict or associated with survival exist, we decided to use a statistical approach which combined PLS and genetic algorithm. Obtaining a gene signature to predict the survival rate is valuable and of practical interest since identification of a set of genes that correlates with the rate of survival can open up avenues for understanding the biological phenomena to explain egg quality and for future biotechnological applications. Our first analysis exhaustively explored the data and provided us with a list of 29 genes that were all considered to be highly related to survival (Table 2). In addition, a second analysis that was performed in order to retain only the genes that were the most pertinent in prediction of developmental competence revealed two solutions that included a set of 7 and 8 genes, respectively (Table 3), that were selected as offering good compromises between quality of the prediction and parsimony of the model genes. There were 5 common genes (italicized) between the two solutions, which could be manipulated for diagnostic purposes. Thus, our statistical modeling approach demonstrated that gene signatures could be obtained from transcriptomic data that could predict developmental competence in fertilized eggs, which would have practical interests.

In an effort to investigate the functional significance of some of the DEGs, we created CRISPR/cas9 knockouts of *otulina* and *sc29a1a* due to their ovary-specific localization, respectively (Fig.5A-5B). Notably, we demonstrated for the first time that deficiency in each of these genes render females subfertile, with complete lack of development in the spawned eggs, which were shown to be unfertilized (Fig. 6r). Thus, our data suggested that *otulina* and *slc29a1a* may play roles that contribute to the factors important for fertilization. *otulina* is predicted to encode for a deubiquitinase, which removes methionine 1-linked ubiquitin chains, of the OTU family in zebrafish, and substrate-bound otulin in mammals has been shown to associate with the linear ubiquitin chain assembly complex (LUBAC). This ubiquitination-deubiquitination system is a key regulator of important signaling pathways, including Wnt, TNF-α, and NF-κb. The *otulin* gene, the mammalian homologue, has been previously shown to be involved in early development in mice since a functionally-disruptive gene mutation results in embryonic lethality due to perturbed Wnt signaling and angiogenesis[17]. In fact, it is known that Wnt signaling plays a major role in gonad differentiation in some fish species[45–47]. Further, *otulin* has also been shown to be a key factor in regulating inflammation and immunity through it’s modulatory role in the TNF-α and NF-κb pathways[18,19]. It is known that inflammatory signaling is an essential part of early embryonic development since many of these components are part of the maternally-inherited repertoire of transcripts, and the TNF-α and NF-κb pathways play important roles in embryonic hematopoietic stem and progenitor cell production as well as body patterning/specification[48–51]. Our results showed that there were significant decreases in the transcript levels of several *wnt* components including *wnt3a, tcf7,lef1*, and *dvl2* (Fig. 7), while none of the transcripts belonging to the *tnf*/*nf*-*κb* pathways showed any changes. Thus, *otulina* deficiency may contribute to subfertility in zebrafish via dysregulation of *wnt* signaling, in line with our previous study that showed that the *wnt* pathway was disturbed at the protein level in bad quality eggs and with the known function of *wnt* in development[31,52,53].

On the other hand, *slc29a1a* is predicted to encode for an equilibrative nucleoside transporter. In mammals, it was shown that *slc29a1* transports adenosine, which is a potent cellular metabolite that functions in cyclic AMP pathways and also acts directly as a vasoactive mediator, into fetal cells and has implications in fetal endothelial functions such that its dysfunction can lead to human pregnancy-related problems such as gestational diabetes, intrauterine growth restriction, and pre-eclampsia[54–56]. In addition, *slc29* homologues in chicken play important roles in rhythm and conduction in developing embryonic hearts via the ERK/MAP (extracellular signal regulated kinase/mitogen activated protein) pathways[57]. However, the function of ***slc29a1*** in fish is still unknown since these species usually undergo external fertilization and embryonic growth. Further investigations into their physiological functions are warranted.

## Conclusions

In this report, transcriptomic profiling of zebrafish fertilized eggs of different quality demonstrated that: 1) dysregulation of the protein synthesis process may be a mechanism behind the reduction in egg quality; 2) gene signatures may exist in the maternally-inherited transcriptome that could be used to predict development competence; and 3) together with the use of CRISPR-cas9 knockout mutants, we clearly showed for the first time that *otulina* and *slc29a1a* are essential for the developmental competence of zebrafish eggs and could be novel maternal-effect genes, which would broaden our understanding of the mechanisms that contribute to egg quality.

## Methods

### Fish husbandry and sample collection

Wildtype zebrafish (*Danio rerio*) of the AB strain were maintained in a central filtration recirculating system with a 12 hr light/dark cycle in the INRA LPGP fish facility (Rennes, France). Individual couple pairing was performed by placing a male and a female overnight in a tank with a partition for separation, and in the morning, the divider was removed after which the female released her eggs to be fertilized by the male. One hundred and thirty-six clutches of fertilized zebrafish eggs at the one-cell stage from 58 families (Additional file 4) were harvested and divided into two parts. One part was flash-frozen in TRI reagent (Sigma-Aldrich, St. Louis, MO) and stored at -80°C for molecular biology analyses. The other part was cultured in modified Yamamoto’s embryo solution (17 mM NaCl, 400 μM KCl, 270 μM CaCl_2_.2H_2_O, 650 μM MgSO_4_.7H_2_O, 0.1 μl/ml of methylene blue) and monitored for up to 48 hours, and the number of survivors was counted at 8, 24, and 48 hpf. Good quality eggs were defined as embryos that had a very high survival rate (>93%) at 48 hpf and bad quality eggs were those that suffered a very low survival rate (<38%) at 48 hpf. All procedures of fish husbandry and sample collection were in accordance with the guidelines set by the French and European regulations on animal welfare. Protocols were approved by the Rennes ethical committee for animal research (CREEA) under approval no. R2012-JB-01.

### RNA extraction

Total RNA of the pooled clutches was extracted using TRI reagent according to the manufacturer’s protocol, and RNA quality and purity were assessed using the Agilent Nano RNA 6000 assay kit and 2100 Bioanalyzer (Agilent Technologies, Santa Clara, CA). All samples were confirmed to have an RIN (RNA integrity number) of 9-10 which are generally accepted as reflecting very good quality RNA.

### Microarray analysis

Zebrafish gene expression profiling was conducted using an Agilent 8×60K high-density oligonucleotide microarray (GPL24500). Labeling and hybridization steps were performed following the Agilent “One-Color Microarray-Based Gene Expression Analysis (Low Input Quick Amp labeling)” protocol. Briefly, for each sample, 150 ng of total RNA was amplified and labeled using Cy3-CTP. Yield (>825 ng cRNA) and specific activity (> 6 pmol of Cy3 per µg of cRNA) of Cy3-cRNA produced were checked with the NanoDrop 2000 spectrophotometer (Thermo Fisher Scientific, Waltham, MA). 600 ng of Cy3-cRNA was fragmented and hybridized on a sub-array. Hybridization was carried out for 17 hours at 65°C in a rotating hybridization oven prior to washing and scanning with an Agilent Scanner (Agilent DNA Microarray Scanner, Agilent Technologies, Massy, France) using the standard parameters for a gene expression 8×60K oligoarray (3 µm and 20 bits). Data were then obtained with the Agilent Feature Extraction software (10.7.3.1) according to the appropriate GE protocol (GE1_107_Sep09) and imported into GeneSpring GX software (Agilent Technologies) for analysis. The data were first normalized by median centering, logged, and then subjected to differential gene expression analysis. All data available in the Gene Expression Omnibus database under accession GSE109073.

### Gene Ontology (GO) analysis

The differentially expressed genes (DEGs) that were found to have a false discovery rate (FDR) < 0.05 and a corrected p-value < 0.05 by microarray were subjected to overrepresentation analyses using DAVID version 6.7 (*https://david.ncifcrf.gov/*)[12] and PANTHER (www.pantherdb.org/)[13] online programs using Ensembl gene identifiers to elucidate enriched terms. The DAVID analyses were conducted using the Functional Annotation Tool based on GOTERM_BP_ALL terms with Benjamini multiple test correction and a FDR<0.05. For the PANTHER analyses, an overrepresentation test, version 10.0 released 2015-05-15, using the GO-Slim Biological Process annotation data set and a Bonferroni correction for multiple testing set at p<0.05 was performed.

### Reverse transcription polymerase chain reaction and quantitative real-time PCR (qPCR)

One μg of RNA was used as template for synthesis of cDNA using the Maxima First Strand cDNA Synthesis Kit (Thermo Fisher Scientific) as per the manufacturer’s protocol. The cDNA samples were then diluted 20-fold and subjected to qPCR using the primers listed in Additional file 2. Primers were designed using the online program Primer3 (http://primer3.ut.ee) and extended across an intron when possible to eliminate the contribution from genomic DNA. qPCR was performed in triplicate using the GoTaq qPCR Mastermix kit (Promega, Madison, WI), which utilizes carboxy-X-rhodamine (CXR) as the reference fluorochrome, using the following cycling condition: 95°C for 10 seconds and 60°C for 30 seconds for 40 cycles. The data were collected with the Applied Biosystems StepOnePlus apparatus (Foster City, CA) and quantitation of the samples was conducted using standard curves. LSM couples member 14B (*lsm14b*), prefoldin subunit 2 (*pfdn2*), and ring finger protein 8 (*rnf8*) had the most stable expression in the microarray dataset and were thus used as internal controls for qPCR. Further, 18S rRNA, *beta-actin* (*bact*), and *elongation factor 1 alpha* (*EF1α*) were also used as internal controls for qPCR[58]. The geometric means of all 6 genes were calculated and for normalization of the data quantity.

### CrispR-cas9 genetic knockout

CRISPR/cas9 guide RNA (gRNA) were designed using the ZiFiT Targeter online software (version 4.2)[59,60] and were made against 3 targets within each gene to generate large genomic deletions, ranging from 130-1500 base pairs, that span exons which allow the formation of non-functional proteins. Nucleotide sequences containing the gRNA were ordered, annealed together, and cloned into the DR274 plasmid.*In vitro* transcription of the gRNA from the T7 initiation site was performed using the Maxiscript T7 kit (Applied Biosystems), and their purity and integrity were assessed using the Agilent RNA 6000 Nano Assay kit and 2100 Bioanalyzer (Agilent Technologies). Zebrafish embryos at the one-cell stage were micro-injected with approximately 30-40 pg of each CRISPR/cas9 guide along with 8-9 nM of purified cas9 protein (a generous gift from Dr. Anne de Cian from the National Museum of Natural History in Paris, France). The embryos were allowed to grow to adulthood, and genotyped using fin clip and PCR that detected the deleted regions. The PCR bands of the mutants were then sent for sequencing to verify the deletion. Once confirmed, the mutant females were mated with wildtype males to produce F1 embryos, whose phenotypes were subsequently recorded. Images were captured with a Nikon AZ100 microscope and DS-Ri1 camera (Tokyo, Japan).

### Genotyping by PCR

Fin clips harvested from animals under anesthesia (0.1% phenoxyethanol) and F1 eggs from females crossed with *vasa:eGFP* males were lysed with 5% chelex containing 100 μg of proteinase K at 55°C for 2 hrs and then 99°C for 10 minutes. The extracted DNA was subjected to PCR using the AccuPrime system (Promega) for *slc29a1a*, Advantage 2 system for *nucleoplasmin 2b* (*npm2b*), and Jumpstart Taq polymerase (Sigma-Aldrich, St. Louis, MO) for *otulina* and *vasa:eGFP*. The primers are listed in Additional file 2.

### Statistical analyses

Statistical analysis of the difference in the expression of each gene between bad and good quality embryos was performed using either Mann-Whitney’s U-test or Student’s t-test after determination of normality of distribution using the Anderson-Darling test. All statistical determinations were conducted using GraphPad Prism version 7 (La Jolla, CA). Data are presented as mean±standard error (SEM). A p-value < 0.05 was considered as statistically significant.

### Analyses by Partial Least Squares (PLS) regression and genetic algorithm

Elimination of the least relevant genes from subsequent analyses was initially performed. First, a correlation test with a 10% p-value threshold between each gene expression and survival rate was conducted. Subsequently, a Pearson correlation coefficient was computed for the expression of each pair of genes. Clusters of genes with pairwise correlations higher than 0.95 were considered as redundant, and only the genes with the highest correlation to survival were selected for subsequent analyses.

Genetic algorithm (GA) was performed whereby *Tpop* (500) random solutions were generated by first choosing a random number (*p*) of genes between 1 and *Pmax* (20) then randomly drawing *p* genes from the candidate genes[61]. PLS regression, which combines the expression levels of the selected genes through linear combinations in order to obtain an estimate of the survival rate, was then applied using the squared Pearson correlation coefficient between the actual survival rates and the estimates as the criterion to evaluate the quality of each proposed subset of genes[62]. A two-fold cross-validation (2-FCV) was always performed to avoid over-fitting, which consisted of applying the PLS model on half of the observations and quantifying the quality of this model by applying it to the other half. This was performed twice by exchanging the application and prediction of the two halves. Once the criterion was computed for each candidate solution, solutions were ranked according to their value, where a high 2-FCV R^2^ reflected a higher rank. Then, selection was applied by associating each solution with a selection probability that was proportional to its quality rank. In this way, the best solutions were selected for subsequent generations whereas suboptimal solutions were likely eliminated[63]. The selected solutions underwent modifications through two operators: mutation and cross-over. Mutation consisted of randomly adding, removing or replacing a gene in the solution. *pm* % (90%) of the solutions underwent mutation. Cross-over consisted of randomly splitting each of two solutions into two subsets of genes and exchanging one subset between them to obtain new gene combinations. *pc* % (50%) of the solutions underwent cross-over. After mutation and cross-over, the new solutions were evaluated again with the same process and selection was applied again. This process was repeated through *Ngene* (200) generations. The solutions selected in the final generation were submitted to extensive evaluation using ten runs of 2-FCV. The average R^2^ obtained for each solution was used as the final criterion to select the best subset of genes.

To challenge the relevancy of the obtained solutions, the same genetic algorithm was applied on randomized datasets, which were obtained by randomly permuting the survival rates of the different observations. First, it is expected that if a gene is linked to the survival process it is likely to often appear in the final solutions. In order to quantify the frequency, the distribution of the number of selections of each gene in the randomized datasets was computed, and only genes with number of selections in the actual data higher than the 95^th^ or 99^th^ percentile of that in the randomized distribution were considered as relevant. Second, the quality of solutions, defined as the distribution of the average 2-FCV R^2^ of the final solutions, obtained on the actual and randomized datasets were also compared. The relationship between gene expression and survival rate was considered significant when the *actual* R^2^ values were significantly higher than the R^2^ values of the randomized dataset. To compare their distributions, a Mann-Whitney test was used.

## List of abbreviations

2-FCV: 2-fold cross validation
*cldn23*: claudin 23
DEGs: differentially expressed genes
*drd3*: dopamine receptor D3
FDR: false discovery rate
*flvcr1*: feline leukemia subgroup C cellular receptor family, member 2a
*gfp*: blue fluorescent protein
hpf: hours post-fertilization
*itih2*: inter-alpha-tryspin inhibitor heavy chain 2
MBT: mid-blastula transition
*npm2a* and *npm2b*: nucleoplasmin 2a/b
*nudt13*: nucleoside diphosphate linked moiety X-type 13
*otulina*: OTU deubiquitinase with linear linkage specificity a, fam105ba
PLS: partial least square regression
*pomt1*: protein-O-mannosyltransferase 1
*prkcq*: protein kinase C, theta
qPCR: quantitative real-time polymerase chain reaction
RNA-seq: RNA sequencing
*rpf2*: ribosome production factor 2 homolog (S. cerevisiae)
*rps27*: ribosomal protein S27 (isoform 2)
*rtn4ip*: reticulon 4 interacting protein 1
*slc29a1a*: solute carrier family 29, member 1a
*spon1b*: spondin 1b
*stra13*: stimulated by retinoic acid 13 homolog/centromere protein X
*tk2*: thymidine kinase 2
*tspan7b*: tetraspanin 7b
*U1*: U1 spliceosomal RNA
WT: wildtype
ZGA: zygotic genome activation

## Declarations

### Ethics approval and consent to participate

Not applicable.

### Consent for publication

Not applicable.

### Availability of data and material

The datasets generated and materials used in the current study are available upon request and in Gene Expression Omnibus (https://www.ncbi.nlm.nih.gov/geo/) database under accession GSE109073.

### Authors contributions

CTC performed most of the experiments and data analyses as well as preparation of the manuscript; TN monitored and harvested the zebrafish embryos and extracted the RNA for microarray; ALC performed the microarray and subsequent data analyses; AP was responsible for animal care and husbandry; LJ facilitated the project through scientific discussions; CR performed the statistical modeling experiment and subsequent data analyses; and JB conceived the study and was responsible for overseeing the project, scientific discussions, and preparation of the manuscript. All authors read and approved the final manuscript.

#### Acknowledgements

We would like to thank all of the support staff and other members of the Fish Physiology and Genomics Institute of Rennes (INRA) for their technical aid. We are very grateful to Jean-Jacques Lareyre (INRA/LPGP) for his kind gift of the Dr_*VaSa*:eGFP C3 zebrafish line.

## Additional Files

Additional file 1: The complete list of the 66 DEGs including the gene description, Ensembl annotation, corrected p-value, and fold change. Analysis was performed with the GeneSpring GX program.

Additional file 2: Sequences of the primer pairs that were used in this study.

Additional file 3: List of differentially expressed genes (DEGs) in bad quality eggs as compared to good quality eggs that were the most modified among the 66 DEGs found by microarray analysis.

Additional file 4: Detailed information on the couples that were mated to produce fertilized eggs and the clutches that were harvested in this study. The number of eggs in each clutch was counted and between 40-60 eggs were followed for up to 48 hours post-fertilization (hpf). The number of surviving embryos were counted at 8, 24, and 48 hpf, and the survival rate was accordingly calculated at each timepoint. * The clutches of bad quality eggs that were used for the couples analysis.

Additional file 5: The complete list of the 1385 DEGs including the gene description, Ensembl annotation, corrected p-value, and fold change from the couples analysis. Analysis was performed with the GeneSpring GX program

Additional file 6: List of 13 differentially expressed genes (DEGs) in bad quality eggs as compared to good quality eggs that were the most modified among the 1385 DEGs found in the couples analysis by microarray.

Additional file 7: Tissue localization of (a) *otulina* and (b) *slc29a1a* transcripts by RNA-seq retrieved from the Phylofish online database.

Additional file 8: Evaluation by qPCR for transcripts of *wnt, tnf*, and *nf-kb* pathways in *otulina* mutant-derived eggs. Transcript levels of (a) *tcf3*, (b) *tnfa*, (c) *ikkaa*, (d), *nf-kb2*, (e) *rel*, and (f) *rela* were investigated by qPCR.

