## Supplementary figures and images for "Lost in translation: egg transcriptome reveals molecular signature to predict developmental success and novel maternal-effect genes"

### Supplementary Materials

**Supplemental Figure 1: Tissue localization of *fam105ba* and *slc29a1a* transcripts by RNA-seq.**

a

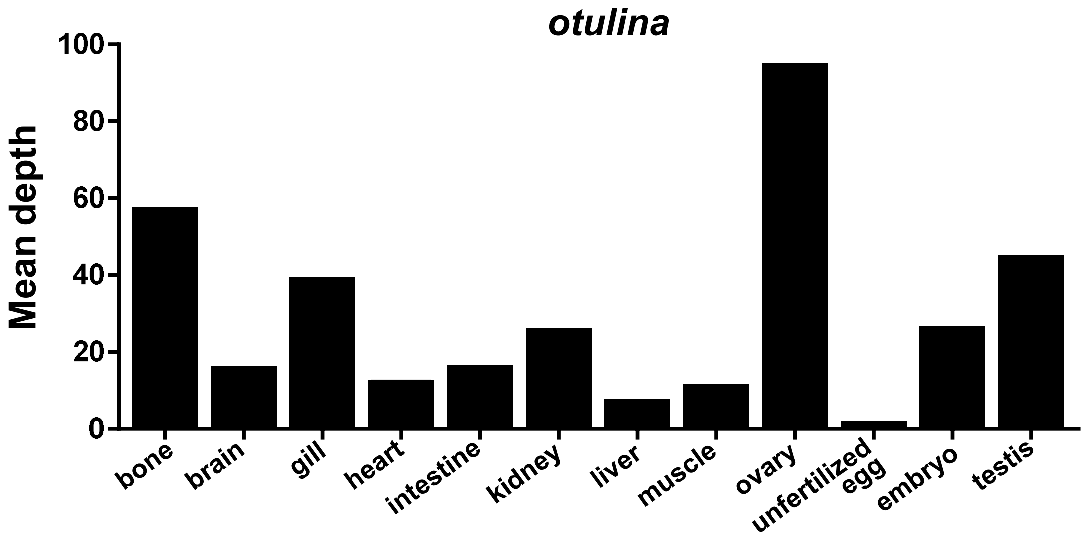


b

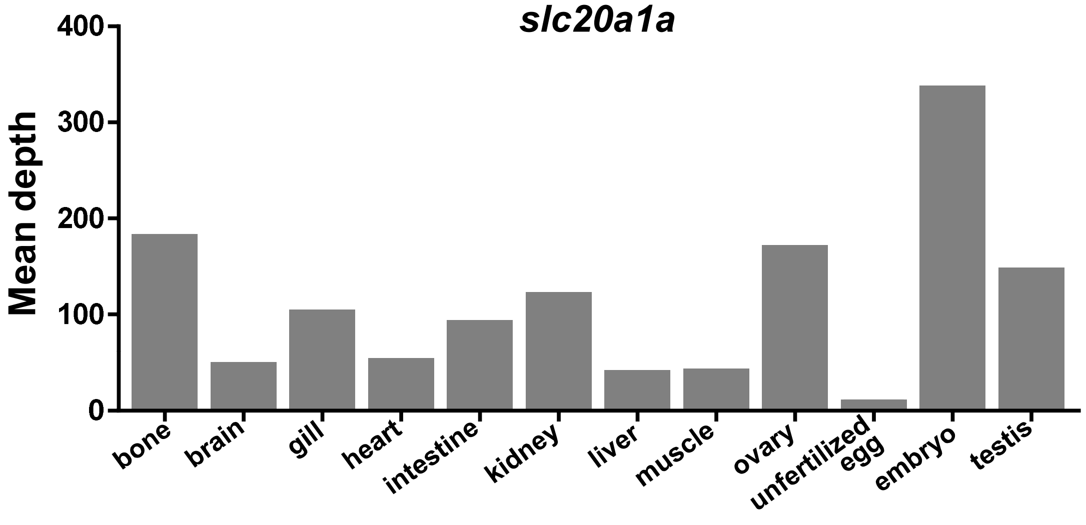
