## Supplementary Materials for "Lost in translation: egg transcriptome reveals molecular signature to predict developmental success and novel maternal-effect genes"

**Supplemental Figure 2: Evaluation by qPCR for transcripts of various pathways in *otulina* mutant-derived eggs.**

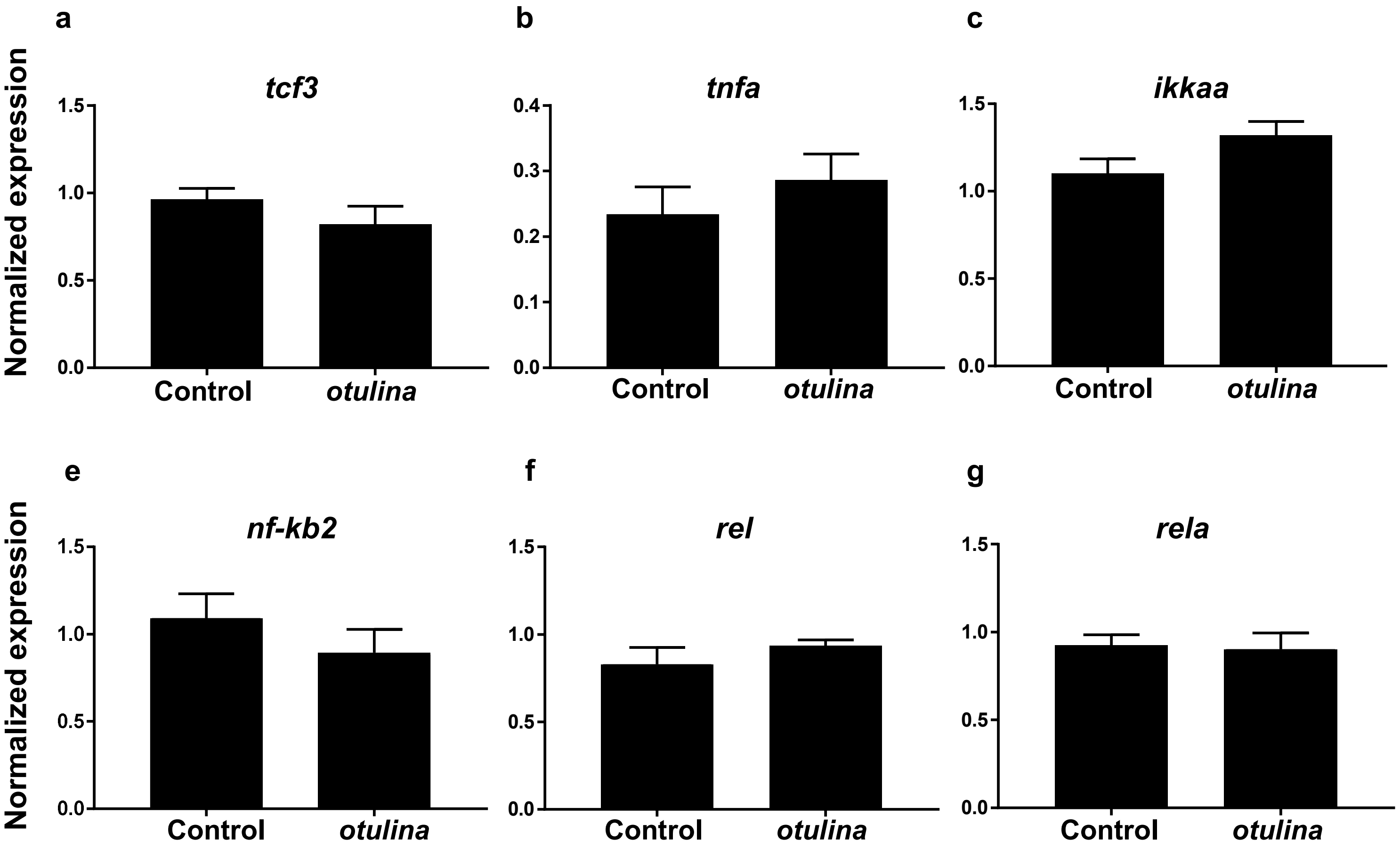
